# Targeting Troponin C with Small Molecules Containing Diphenyl Moieties: Calcium Sensitivity Effects on Striated Muscle and Structure Activity Relationship

**DOI:** 10.1101/2023.02.06.527323

**Authors:** Eric R. Hantz, Svetlana B. Tikunova, Natalya Belevych, Jonathan P. Davis, Peter J. Reiser, Steffen Lindert

## Abstract

Despite large investments from academia and industry, heart failure, which results from a disruption of the contractile apparatus, remains a leading cause of death. Cardiac muscle contraction is a calcium-dependent mechanism, which is regulated by the troponin protein complex (cTn) and specifically by the N-terminal domain of its calcium binding subunit (cNTnC). There is an increasing need for the development of small molecules that increase calcium sensitivity without altering systolic calcium concentration, thereby strengthening cardiac function. Here, we examined the effect of our previously identified calcium sensitizing small molecule, ChemBridge compound 7930079, in the context of several homologous muscle systems. The effect of this molecule on force generation in isolated cardiac trabeculae and slow skeletal muscle fibers was measured. Furthermore, we explored the use of Gaussian accelerated molecular dynamics in sampling highly predictive receptor conformations based on NMR derived starting structures. Additionally, we took a rational computational approach for lead optimization based on lipophilic diphenyl moieties. This led to the identification of three novel low affinity binders, which had similar binding affinities to known positive inotrope trifluoperazine. The most potent identified calcium sensitizer was compound 16 with an apparent affinity of 117 ± 17 *μM*.

## Introduction

The occurrence of heart failure has been shown to increase over time with aging of the population, and its impact was estimated at $216 billion in direct cost from 2016 - 2017 ^1^. Heart failure (HF) has been generally characterized as the heart’s inability to pump and/or fill with enough blood to meet the demands of circulation ^2^. Heart failure can be divided into three subtypes: HF with reduced ejection fraction (HFrEF), heart failure with preserved ejection fraction (HFpEF), and HF with mid-range ejection fraction (HFmrEF). These subtypes have been differentiated based on the percentage of ejection fraction (EF) from the left ventricle, where HFrEF has an EF below 40%, HFpEF has an EF greater than 50%, and HFmrEF has an EF ranging from 40-50% ^3, 4^. HF has been classically treated with diuretics and blood pressure medication to reduce blood volume. However, the focus for many experts is the design of small molecules that restore cardiac contractility and functionality in efforts to treat HFrEF. A popular class of small molecules with this intended purpose are positive inotropes ^5, 6, 7, 8, 9^. Some of these compounds have been shown to utilize the beta-adrenergic pathways to create an increased systole concentration of Ca^2+^ ions. However, increasing the systolic [Ca^2+^] can potentially lead to arrhythmias in patients ^10, 11^. The current direction of the field has shifted towards the identification and development of positive inotropes that increase Ca^2+^ sensitivity of the proteins involved in muscle contraction without affecting the concentration of systolic Ca^2+^ ^12, 13, 14, 15, 16, 17, 18, 19, 20, 21, 22^.

Ca^2+^-dependent heart muscle contraction is regulated by cardiac troponin (cTn). The cTn complex consists of three subunits: troponin T (cTnT) which interacts with tropomyosin and anchors the protein complex to the thin filament, troponin I (cTnI) known as the inhibitory subunit, and troponin C (cTnC) which facilitates Ca^2+^ binding. cTnI plays a key structural role in anchoring cTnC to the rest of the troponin complex, in addition to having two amphiphilic peptides that play important functional roles. Under unsaturated Ca^2+^ conditions, the cTnI inhibitory peptide (residues 137-148) interacts with actin, thereby inhibiting actomyosin binding ^23^. Additionally, the cTnI switch peptide (cTnI_sp_ residues 149-164) interacts with the N-terminal domain of cTnC stabilizing the open conformation of the domain when saturated with Ca^2+^ ^23, 24, 25^. cTnC is a dumbbell-shaped protein with two globular domains connected by a flexible linker, each containing two EF-hand motifs ^24, 26^. The C-terminal domain (cCTnC), termed the structural domain, contains two high-affinity binding sites (sites III and IV) which are constantly saturated by Ca^2+^ or Mg^2+^ under physiological conditions ^27^. The N-terminal domain (cNTnC), known as the regulatory domain, contains only one active binding site (site II) ^28, 29^. Upon Ca^2+^ binding to site II, a large conformational change of the troponin complex is initiated by cNTnC. One feature of this structural rearrangement is the opening of a hydrophobic patch (residues 20, 23, 24, 26, 27, 36, 41, 44, 48, 57, 60, 77, 80, and 81) in cNTnC, which becomes stabilized upon binding of the cTnI_sp_. The hydrophobic patch serves as an attractive target for increasing Ca^2+^ sensitivity via small molecules. Additionally, the hydrophobic patch and entire troponin complex have been the focus of numerous in silico studies to capture the proteins dynamics of hydrophobic patch opening^30, 31, 32, 33, 34, 35, 36, 37^, and cTnI_sp_ binding ^38, 39, 40, 41, 42^.

We have knowledge of several compounds that bind to the hydrophobic patch and modulate the sensitivity of Ca^2+^ binding: levosimendan ^43^, one of the most well-known Ca^2+^ sensitizers, and its analog pimobendan ^44^, 3-methyldiphenylamine (3-mDPA) ^45^, bepridil ^46^, W7 ^47, 48^, dfbp-o (2′,4′-difluoro(1,1′-biphenyl)-4-yloxy acetic acid) ^49^, trifluoperazine (TFP) ^50^, National Cancer Institute (NCI) database compounds NSC147866 ^18^, NSC600285 ^16^, NSC611817 ^16^, and ChemBridge compounds 6872062, 7930079, and 9008625 ^17^. The two-dimensional structures of these compounds are shown in **Figure 1**. There are currently no FDA approved Ca^2+^ sensitizers, despite several of these compounds proceeding to clinical trial. Levosimendan has been utilized in Europe for decades ^51^, but has yet to receive FDA approval. Compounds trifluoperazine and bepridil failed clinical trials due to off-target effects ^52^. This underscores the need for continual improvement in positive inotrope efficacy and safety.

**Figure 1.**
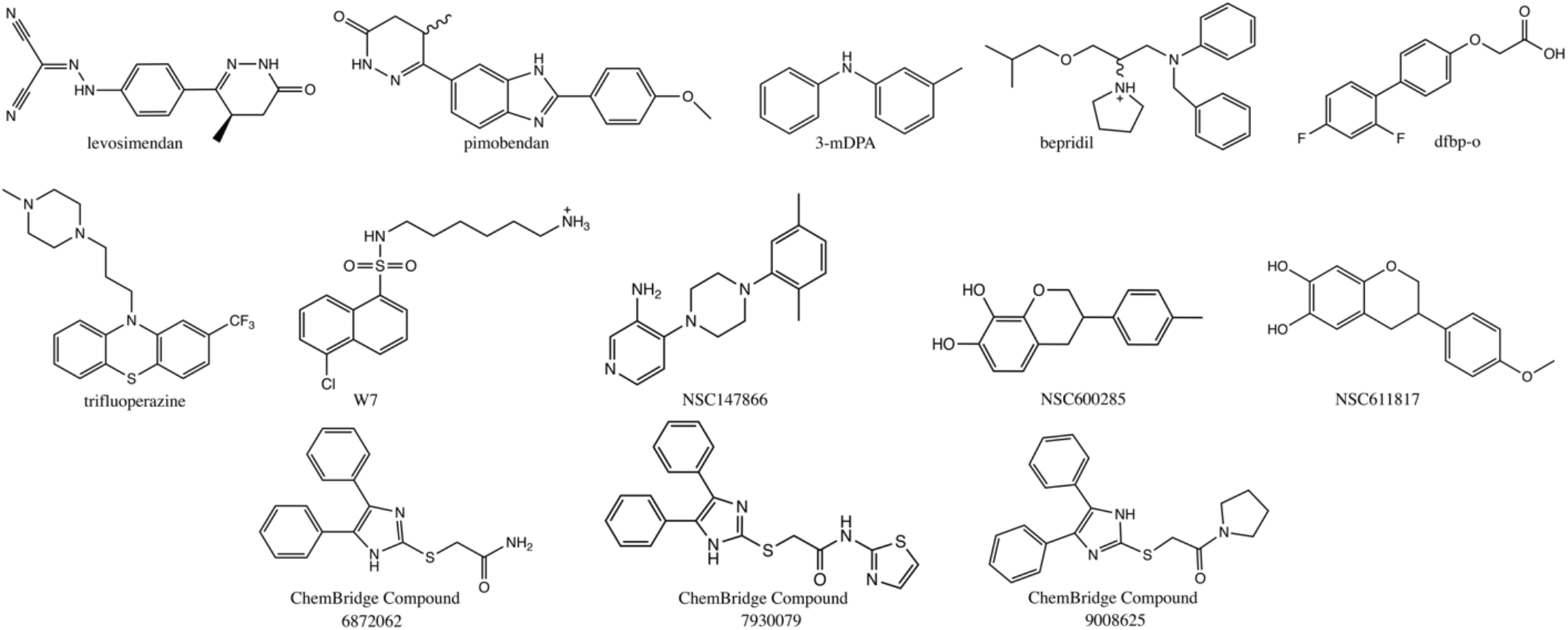
2D structure of known cNTnC Ca^2+^ sensitivity modulators.

In previous work we identified three Ca^2+^ sensitizing compounds with respective *K*_*D*_ values < 100 *μM*, with the lead compound being ChemBridge compound 7930079 (2-[(4,5-diphenyl-1H-imidazol-2-yl)thio]-N-1,3-thiazol-2-ylacetamide). This compound is one of the most potent and high affinity cNTnC binders known to date. In this work, we examined the sensitizing effects of 2-[(4,5-diphenyl-1H-imidazol-2-yl)thio]-N-1,3-thiazol-2-ylacetamide in rat cardiac trabeculae and in slow skeletal muscle fibers. We found this compound induced a small but significant shift in cardiac trabeculae and a much larger (four-fold) shift in slow skeletal fibers. Furthermore, we evaluated the use of Gaussian accelerated Molecular Dynamics (GaMD) in generating predictive receptor conformations for virtual screening based on initial NMR model starting structures. Additionally, we further explored the structure activity relationship of potential Ca^2+^ sensitizers based on the lipophilic character of the known high affinity Ca^2+^ sensitivity modulators. We found three compounds to slow the rate of Ca^2+^ dissociation from cNTnC within a cTnC-cTnI chimera, with apparent disassociation constants ranging from 117 − 482 *μM*, similar to that of many known Ca^2+^ modulators.

## Methods

### Receptor Selection and Preparation

For our computational docking studies, we seek to identify cNTnC – cTnI_sp_ conformations that are highly predictive. We prioritized the inclusion of the cTnI_sp_ to accurately model the cTnC-cTnI chimera utilized in our biochemical studies. Thus, we performed an exhaustive search for human cNTnC – cTnI_sp_ structures with known Ca^2+^ sensitivity modulating small molecules bound in the hydrophobic patch (holo) and ligand unbound (apo) structures in the RCSB protein data bank. We obtained apo structures from two PDB entries: 1MXL ^53^ where cNTnC is in complex with the cardiac isoform of the TnI switch peptide and 2MKP ^54^ where cNTnC is in complex with the fast skeletal TnI switch peptide. We obtained holo cNTnC conformations of several known calcium sensitivity modulators: bepridil (PDB: 1LXF ^55^), W7 (PDBs: 2KFX ^56^, 2KRD ^57^, and 6MV3 ^47^), dfbp-o (PDB: 2L1R ^49^), and 3-mDPA (PDBs: 5WCL ^45^ and 5W88 ^45^). All selected PDB entries were obtained from NMR experiments and contained multiple conformers; each structure was extracted resulting in 166 receptor conformations. We examined all 166 receptor conformers in our docking studies.

All receptor conformations were imported into Schrödinger’s Maestro ^58^ and prepared using the Protein Preparation Wizard ^59^; where for all applicable conformers the C-terminus was capped by the addition of an *N*-methyl amide and the N-terminus was capped by the addition of an acetyl group. The protonation states of all titratable residues were assigned using EPIK ^60, 61^ with a pH constraint of 7.4 ± 1.0.

### GaMD Simulations and Clustered Receptor Generation

In previous work we have demonstrated the utility of GaMD to generate highly predictive receptor conformations based on X-ray crystallographic structures ^62^. Here, we evaluated the impact of this computational technique for increased sampling of troponin’s structural ensemble when the initial frame was derived from an NMR experiment. For each of the beforementioned nine NMR PDB deposited structures (1MXL, 2MKP, 1LXF, 2KFX, 2KRD, 6MV3, 2L1R, 5WCL, and 5W88), we extracted model one (the representative receptor conformer) which served as the initial starting frame for a 300 ns GaMD simulation performed with Amber20 ^63, 64^. For ligand-bound conformers, the small molecule was parameterized using the second generation of the generalized amber forcefield (GAFF2) ^65^ and Amber’s antechamber software. All protein complexes were parameterized with the amber forcefield ff14SB ^66^, solvated with TIP3P ^67^ water molecules in a 12 Å octahedron, and neutralized with sodium ions.

All systems were minimized with restraints (20 kcal/mol) on the protein and ligand (if applicable) using 2500 steps of steepest decent minimization followed by 2500 steps of conjugate gradient decent. A second unconstrained minimization was sequentially performed using 2500 steps of steepest decent minimization followed by 2500 steps of conjugate gradient decent. The system was then heated to 310 K over a span of 1 ns using the Langevin thermostat ^68^. For each system, a short conventional MD simulation was performed for equilibration under constant temperature (310 K) and pressure (1 bar) using the Langevin thermostat and Berendsen barostat ^69^, prior to the GaMD preparation simulations. During the GaMD preparation simulation statistics were collected to calculate the appropriate boosts to apply to the dihedral and total potential energies. These statistics were obtained from a second 10 ns conventional MD run, the initial boosts then applied, and subsequently updated during a 50 ns GaMD biasing run. The final GaMD restart parameters (VmaxP, VminP, VavgP, sigmaVP, VmaxD, VminD, VavgD, and sigmaVD) were then read in for a 300 ns GaMD production run. The upper limit for the dihedral and total boost potentials was set to 6 kcal/mol. All simulations were performed with a 12 Å cutoff for electrostatic and van der Waals interactions, and a 2 fs timestep with the SHAKE algorithm ^70^. Coordinates were saved every 2 ps, resulting in 150,000 frames.

The final structures used for docking studies were obtained by clustering each of the 300 ns GaMD simulations individually. Prior to clustering the individual trajectories, all water molecules, ions, and ligands were removed. Clustering was performed over every fourth frame of the original 150,000 frames, resulting in 37,500 frames available for clustering. The density-based clustering algorithm (DBScan) ^71^ implemented in Amber’s CPPTRAJ was used to cluster the processed trajectories to obtain approximately 11 representative conformers per simulation. Trajectories were clustered using the backbone atoms of the hydrophobic patch residues (residues 20, 23, 24, 26, 27, 36, 41, 44, 48, 57, 60, 77, 80, and 81). In total, 101 clustered conformers were generated and used in further active/decoy docking studies (see below). All receptor conformers were loaded into Maestro and prepared using the Protein Preparation Wizard tool, as detailed above.

### Receptor Grid Generation

Receptor grids for ligand-bound receptor conformers were generated by selecting the ligand within the Maestro workspace and using the center of mass of the ligand as the center of the receptor grid. For apo receptor conformers, the center of the search space was determined by submitting a PDB file of the conformer to the FTMap webserver ^72, 73^, where fragments were globally docked to identify potential small molecule binding sites. The resulting output file (PDB format) was imported into PyMOL ^74^, where the align function was utilized to overlay the FTMap PDB file with a holo receptor conformer. Upon visual inspection, the fragment cluster that overlapped the position of the known calcium modulator was identified, and the center of mass of the cluster was calculated. The center of mass served as the three-dimensional coordinates for the center of the receptor grid. The search area was centered on the ligand’s center of mass or the supplied three-dimensional coordinates, respectively, and allowed the centroids of any docked compound to fully explore a 10 × 10 × 10 Å^”^ inner search space, while the periphery of the ligand was able to extend out into a 20 × 20 × 20 Å^”^ search space. The OPLS3e forcefield ^75^ was used to generate the search grid and all hydroxyl groups were selected to be freely rotatable within the search area.

### Ligand Preparation

The three-dimensional coordinates of all small molecules (known binders) were extracted from the representative conformation of their respective PDB entries. Small molecules from the Schrödinger decoy sets, the ChemBridge EXPRESS-Pick Collection, the ChemBridge Core Library, and NCI database used in our docking protocol were obtained from their respective databases in the form of SDF files containing their three-dimensional structural information. For the known binders levosimendan, pimobendan, and trifluoperazine, we generated their respective structures using Schrödinger’s 2D Sketcher tool. Schrödinger’s LipPrep^76^ tool was used to prepare each ligand for docking. All tautomers, protomers, and stereoisomers were generated for each ligand. Protonation states were assigned using EPIK with a pH value of 7.4 ± 1.0, identical to that of the protein preparation step.

### Active/Decoy Screening and Receptor Performance Analysis

A total of 267 receptor conformations (166 NMR receptor conformers and 101 GaMD clustered conformers) were evaluated by active/decoy screening. The 13 known sensitivity modulators were used as known actives in the active/decoy docking process, and the presumed decoy small molecules (1,000) were obtained from the Schrödinger decoy set ^77^ with an averaged molecular weight of 360 g/mol. The decoy set was confirmed to match the molecular properties of the known binders based on the following chemical properties: molecular weight, LogP, formal charge, number of hydrogen bond donors, and number of hydrogen bond acceptors. These properties were calculated using the RDKit (version 2020.09.1) ^78^ Chem Descriptor ExactMolWt, Crippen model, GetFormalCharge, Lipinski NOCount, and Lipinski NHOHCount modules, respectively (see **Figure S1**). All compounds were subject to ligand preparation as detailed above. The active and decoys compounds post-LigPrep were docked into all receptor conformations using Schrödinger’s Glide SP ^77, 79, 80^. Default parameters from the Schrödinger 2018-3 release were utilized for all docking simulations.

The docked poses of all small molecules were ranked by their respective docking score. Subsequently, the top docked pose of each compound was kept, and all other protomers/tautomers/stereoisomers of the compound removed. The performance of every receptor conformer was evaluated with three metrics: receiver operating characteristic (ROC) curve, enrichment factor, and a weighted enrichment factor. The ROC curve was generated by plotting the true positive rate (TPR) against the false positive rate (FPR) and the area under the ROC curve (AUC) was calculated using Python’s scikit-learn library (ver.0.22.1) ^81^. The ROC AUC metric evaluates the predictiveness of the receptor conformation for use in future blind screenings. Enrichment factor (EF) was calculated to evaluate the number of known binders for a predefined early recognition period (in our case: top 40 compounds). The early recognition period was determined based on the number of compounds we typically screen in one iteration of our *in vitro* studies. EF was calculated via the following equation:

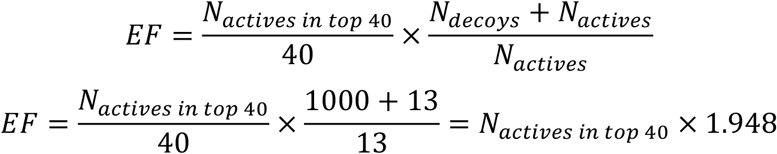

For cNTnC, nine of the thirteen known actives were classified as poor binders (*K*_*D*_ > 100*μM*), and four actives (ChemBridge compounds 6872062, 7930079, 9008625, and 3-mDPA) were classified as high – moderate affinity binders (*K*_*D*_ < 100*μM*). We aimed to minimize the influence of the poor binders with regards to the EF score and subsequent conformer identification, as we believed this would steer the computational screenings towards the identification of additional poor binders. Therefore, in order to prioritize the identification of the high affinity known binders in the early recognition period, we formulated a weighted enrichment factor (*EF*_*weighted*_) score:

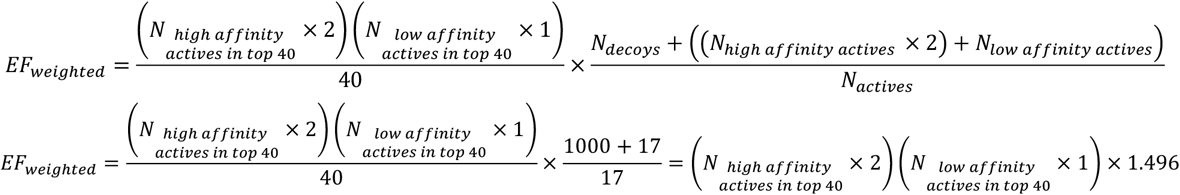

### Computational Investigation of Compounds with Diphenyl Groups

Based on the two-dimensional structure of the four known high affinity Ca^2+^ sensitizers (3-mDPA and ChemBridge compounds 6872062, 7930079, and 9008625), we hypothesized that the diphenyl rings are crucial for ligand binding. In order to preserve the lipophilic interactions, we created three chemical motifs (diphenyl-imidazole, diphenyl-triazine, and diphenylmethane; see **Figure S2**) for two-dimensional structural similarity searches of large chemical libraries. In preparation for our database searches, we filtered both the ChemBridge EXPRESS-Pick Library (501,916 compounds) and Core Library (812,681 compounds) with a cheminformatics approach. Similarly to our previous work ^17^, compounds found to have a molecular weight greater than 400 g/mol or a calculated LogP value less than 2.0 or greater than 4.0 were removed. In a second filtering step, compounds found to violate both of the following rules were removed as violators: number of hydrogen bond donors > 5 and number of hydrogen bond acceptors > 10. Additionally, the libraries were screened for Pan-assay interference (PAINS) compounds and violators were sequentially removed. Upon completion of library filtering, the EXPRESS-Pick library was reduced to 206,726 compounds and the Core library was reduced to 412,866 compounds. For each of the remaining compounds (619,592 compounds), we calculated the Tanimoto coefficient between that compound and the three diphenyl motifs. We identified all compounds that had a Tanimoto coefficient *≥* 0.6 to any of the diphenyl functional groups, resulting in 89 compounds. The compounds were further filtered by removing any compounds we had tested in our previous study ^17^ to avoid overlap, compounds that did not contain a diphenyl motif via visual confirmation, and any compounds that added functional groups exclusively to the diphenyl rings. The last filtering step was implemented to prioritize identification of compounds with extensions from the imidazole, triazine, or methane in efforts to retain similar protein-ligand interactions as compared to the thiazole acetamide tail of 2-[(4,5-diphenyl-1H-imidazol-2-yl)thio]-N-1,3-thiazol-2-ylacetamide (see **Figure S3**). These criteria further reduced the number of compounds to 36 compounds. All remaining 36 compounds were docked into the top three performing receptor conformations (1LXF M6, 1LXF M23, and 1LXF M28) based on the results from our active/decoy docking simulations. The compounds were all subjected to ligand preparation as detailed above, and docked using Glide SP with default settings, and the docking score was averaged across the three conformers to select a top set of potentially promising compounds for experimental testing. Compounds that exhibited an averaged docked score across the three conformers of > −5.0 *kcal/mol* were removed from experimental testing consideration. This resulted in 23 compounds to be ordered for *in vitro* testing based on Tanimoto coefficients.

In addition to identifying potential sensitizing compounds based on two-dimensional similarity, we performed an exhaustive blind docking of the ChemBridge Core Library. We utilized the filtered version of the Core Library as described above. The 412,866 compounds were prepared as described in the Ligand Preparation methods section, and subsequentially docked into the top three receptor conformations (1LXF M6, 1LXF M23, and 1LXF M28) using Glide SP with default settings. Upon completion of docking, the docking Z-score of each compound in each receptor was calculated and averaged over all conformers. We have previously shown that compound selection with an unbiased ranking such as Z-score, can lead to high success rates in blind virtual screenings ^62, 82^. Based on ranking by averaged Z-score, the two-dimensional structures of the top 40 compounds were visually inspected and diphenyl motifs were identified in 17 of the compounds. We selected these 17 diphenyl containing compounds for experimental testing. In total, 40 compounds (23 based on Tanimoto coefficients and 17 from the Core library blind screening) were ordered from ChemBridge and tested *in vitro*.

### Preparation of Proteins for Biochemical Studies

The cTnC-cTnI chimera (herein designated as chimera) was generated as previously described ^17,83^. The purified chimera was labeled with the environmentally sensitive fluorescent probe IAANS on Cys53 of the cNTnC (within the chimera) with C35S, T53C and C84S substitutions, as previously described ^84^

### Stopped-Flow Fluorescence Measurements

All kinetic measurements were carried out in stopped-flow buffer (10 mM MOPS, 150 mM KCl, pH 7.0) at 15°C using an Applied Photophysics Ltd. (Leatherhead, UK) model SX.18MV stopped-flow apparatus with a dead time of ∼1.4 ms. IAANS fluorescence was excited at 330 nm and monitored using a 420-470 nm band-pass interference filter (Oriel, Stratford, CT). Ca^2+^ chelator EGTA (10 mM) in stopped-flow buffer was used to remove Ca^2+^ (200 μM) from the chimera (0.5 μM) also in stopped-flow buffer in the absence or presence of compounds. Varying concentration of each compound were individually added to both stopped-flow reactants. Data traces were fit using a program by P.J. King (Applied Photophysics, Ltd.), that utilizes the non-linear Lavenberg-Marquardt algorithm. Each k_off_ represents an average of at least three separate experiments ± standard error, each averaging at least three shots fit with a single exponential equation.

### Calcium-sensitivity of Force Generation

The care and use of the animals (four male Sprague Dawley rats, 7-9 months old) used for this project were approved by the Institutional Animal Care and Use Committee of Ohio State University. The preparation of cardiac trabeculae (n=10) and slow skeletal muscle (soleus) fibers (n=10) was as described in Tikunova et al ^13^. The force measurements were also as described in Tikunova et al ^13^. Briefly, active force generation was measured in a series of activating solutions with different pCa (-log of [Ca^2+^]) values, first without, then with the Compound 7930079 (20 μM). The force versus pCa data were fit as described in Mahmud et al. 2022 ^15^. The pCa that corresponded to 50% of maximal force generation (pCa_50_) was determined from the curve fits and the difference in the value between with and without the compound was calculated for each cardiac trabecula and muscle fiber. The paired Student’s t-test was used to evaluate the statistical significance of the difference in pCa_50_ with and without the compound. The fiber type of the soleus fibers that were used for the force/pCa measurements was determined with SDS-PAGE, as described in Bergrin et al., 2006 ^85^.

## Results and Discussion

### Effect of Compound 7930079 on Force Generation in Isolated Cardiac Trabeculae and Slow Skeletal Muscle Fibers

All of the soleus fibers that were selected for force measurements were verified to be slow-type, based upon the results from SDS-PAGE (**Figure S4**). A summary of the force/pCa data obtained in cardiac trabeculae and in skeletal slow fibers, with and then without 20 *μ*M 7930079, is shown in **Figure 2**. This compound significantly (P<.01) shifted the curve to higher sensitivity in cardiac trabeculae (0.05 ± 0.04 pCa unit) and in slow fibers (0.20 ± 0.03 pCa unit). We also tested whether 7930079 impacted maximal force generation in cardiac trabeculae and in slow fibers. Every third activation in the series of force/pCa measurements was in pCa 4.0 activating solution which yields maximal force generation in both types of preparation. The peak force generated in pCa 4.0 solution was averaged (n=6 activations), without then with, the compound. Compound 7930079 had no effect (P=0.39) on average maximal force generation in slow fibers. There was a significant (P=0.001) decrease of 8% in the average force in the presence of 7930079 in cardiac trabeculae. However, in a control set of measurements in six other cardiac trabeculae, in which the complete force/pCa relationship was measured twice with the compound, the average maximal force in pCa 4.0 activating solution was 6%, lower, on average, in the second, compared to the first, series of measurements in each trabecula. We, therefore, conclude that 7930093 caused a very small (∼2%) reduction in maximal force generation in cardiac trabeculae. While these are promising results, this underscores the need for further identification of Ca^2+^ sensitizers and of particular interest in this study, the exploration of structure-activity relationships around the diphenyl motifs.

**Figure 2.**
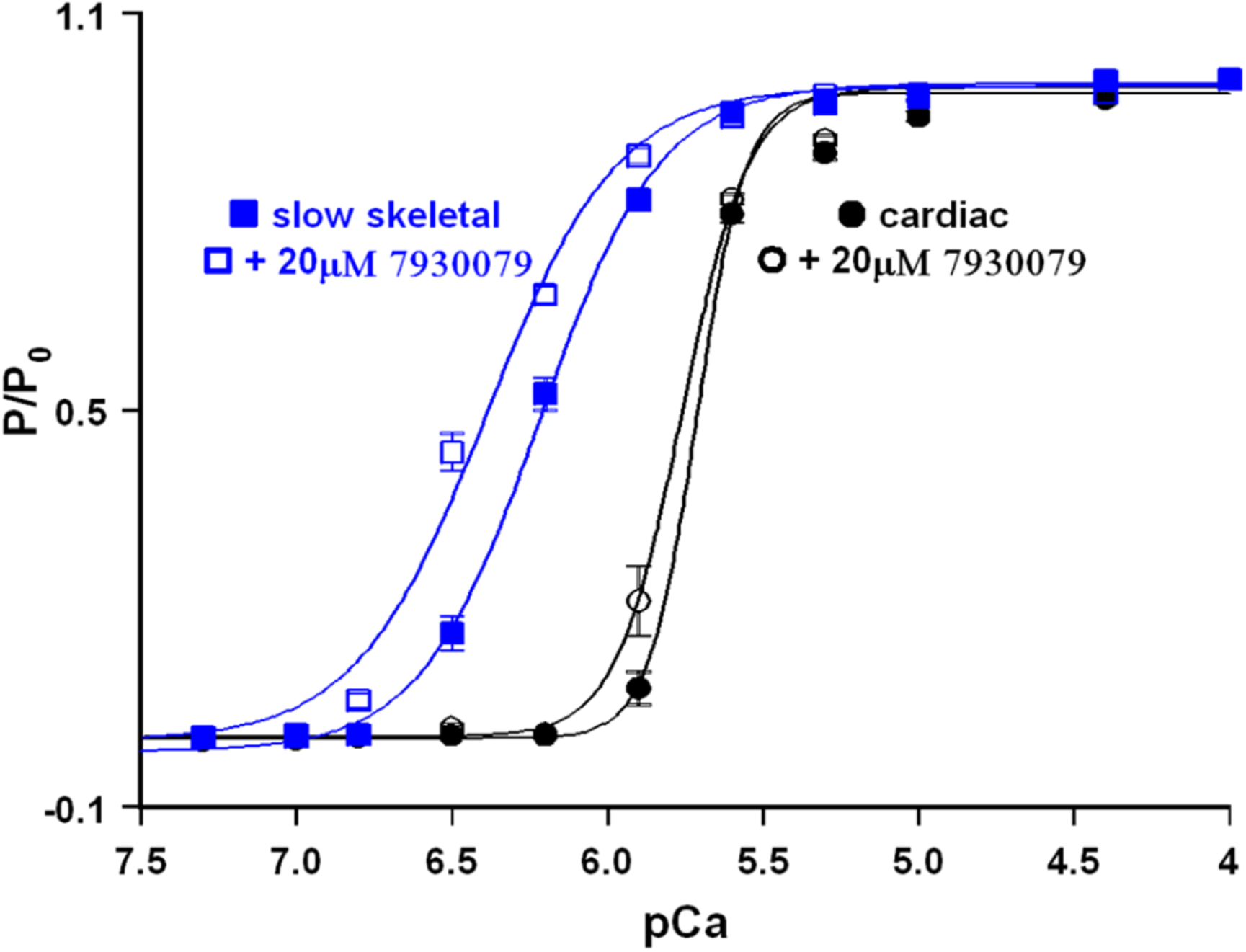
The Force/pCa relationships were measured in both rat cardiac trabeculae and slow skeletal rat muscle fibers (10 each) before and after the addition of compound 7930079, our highest affinity compound to date. Compound 7930079 significantly (P<0.05) sensitized both cardiac, and more potently, slow skeletal muscle to Ca^2+^, as evidenced by the increase in Ca^2+^ sensitivity (positive ΔpCa50) compared to the absence of the compound (internal control).

### Top Receptor Conformations Identified Based on Active/Decoy Docking

Prior to virtual screens of small molecule libraries, we sought to further characterize the predictiveness of PDB-deposited cNTnC-cTnI_sp_ receptor conformations to correctly identify known cNTnC binders. We identified 166 NMR-derived conformers for characterization, expanding our search from previous works ^16,17^. Additionally, we sought to determine the effectiveness of Gaussian accelerated Molecular Dynamics (GaMD) generated conformers based on NMR starting structures. Recently, we have shown that clustering over GaMD simulations may result in the identification of additional highly predictive receptor conformations ^62^. However, the starting structures for this study were almost exclusively from x-ray crystallography structures deposited in the PDB. Here, for the cNTnC-cTnI_sp_ system, we aimed to determine if this enhanced sampling technique could capture additional protein dynamics that are currently absent from known NMR PDB structures. Therefore, based on the representative NMR model of each PDB entry (as described in Methods), we performed a 300 ns GaMD simulation and clustered over each individual trajectory, resulting in 101 additional receptor conformations.

In total 267 cNTnC-cTnI_sp_ conformers were used in our active/decoy screening, where all compounds were docked utilizing the Glide SP docking methodology. The 13 known cNTnC Ca^2+^ sensitivity modulators served as the known actives and were docked along a set of 1,000 presumed decoy small molecules. The decoy compounds were obtained from Schrödinger’s decoy set with an average molecular weight of 360 g/mol. In order to avoid any unrealistic enrichment in the active/decoy screenings, the decoys and actives were property-matched across five chemical properties: molecular weight, LogP, formal charge, number of hydrogen bond acceptors and number of hydrogen bond donors (see **Figure S1**). For all conformers, we calculated the respective true positive rates (TPRs) and false positive rates (FPRs) and determined the AUC of the generated ROC curves. The ROC AUC metric was used to evaluate the predictiveness of the conformer in identifying known actives over the decoy compounds. Furthermore, we used enrichment factor (EF) as a confidence metric for identifying the actives in a predefined early recognition period (top 40 compounds). However, particularly in the case of cTnC, many of the known actives can be classified as weak binders (*K*_*D*_ > 100 *μM*) to the hydrophobic patch of cNTnC in the presence of cTnI_sp_, such as: levosimendan (*K*_*D*_ ≈ 700 *μM*) ^86^, bepridil (*K*_*D*_ = 380 ± 60 *μM*) ^13^, W7 (*K*_*D*_ = 150 − 300 *μM*) ^87^, dfbp-o (*K*_*D*_ = 270 ± 30 *μM*) ^49^, TFP (*K*_*D*_ = 110 ± 20 *μM*) ^13^, and NCI compound NSC147866 (*K*_*D*_ = 379 ± 50 *μM*) ^18^. The experimental binding affinities of the other two NCI compounds (NSC600285 and NSC611817) were not reported but were predicted to be weak binders. In order to dampen the influence of the weak binders for the purpose of receptor selection, we developed a weighted enrichment factor (EF_weighted_) score to prioritize the early identification of known cNTnC Ca^2+^ sensitivity modulators with moderate to high affinity. Those included 3-mDPA (*K*_*D*_ = 30 *μM*) ^45^ and ChemBridge compounds 6872062, 7930079, and 9008625 with respective apparent experimental binding affinities of 84 ± 30 *μM*, 1.45 ± 0.09 *μM*, and 34 ± 12 *μM* ^17^. These four compounds contain diphenyl motifs, which served as the inspiration for our two-dimensional similarity searches of the ChemBridge databases (see below). The data for all three metrics (ROC AUC, EF, and EF_weighted_) for all 166 NMR conformers is summarized in **Table S1**, and the data for all 101 clustered GaMD conformers is summarized in **Table S2**.

**Figure 3** shows the evaluation of all 267 receptor conformations, as well as the ROC curve of the top performing receptor conformation (1LXF NMR model 6) with the respective values for ROC AUC, EF, and EF_weighted_ explicitly shown. The top three receptor conformations were determined to be all NMR models of PDB entry 1LXF (1LXF model 6, 1LXF model 23, and 1LXF model 28), and were used in later virtual screens. Interestingly, the top performing conformers all belong to the same PDB (1LXF), which contains cNTnC in complex with cTnI_sp_ and bepridil. We believe that our focus on the diphenyl motifs (through use of the weighted EF) led to implicit bias of selecting 1LXF receptor conformations. 1LXF is the only deposited structure of the cNTnC – cTnI_sp_ complexes with a small molecule containing a diphenyl motif (benzyl methylaniline). Therefore, we speculated, the various receptor conformations belonging to this PDB entry could capture more favorable protein-ligand interactions between the hydrophobic patch residues and that of compounds with a diphenyl structure. Additionally, only one GaMD clustered conformer (5WCL clustered conformer 6) exhibited a relatively high ROC AUC of 0.750; however, the EF and EF_weighted_ values were low (5.89 and 5.98 respectively). As shown before, GaMD is a powerful technique for structure-based drug discovery, with the most benefit seen when x-ray crystallographic structures served as the initial frame of the simulation. Unfortunately, in our case, there was no improvement for capturing ideal receptor conformations for ligand binding compared to preexisting NMR based structures. The top three receptor conformers (1LXF model 6, 1LXF model 23, and 1LXF model 28) were used for all virtual screenings going forward.

**Figure 3.**
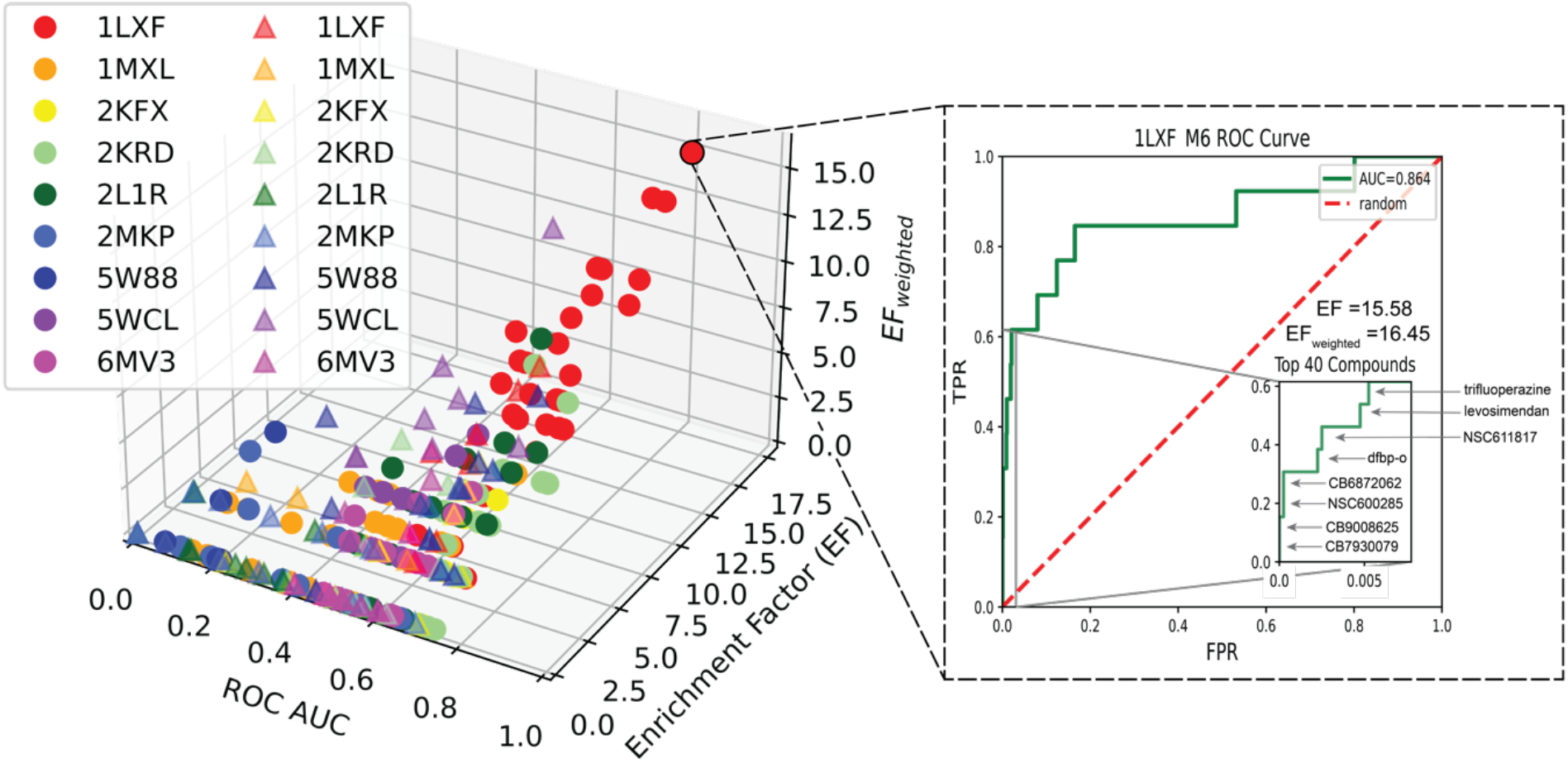
cNTnC receptor conformation characterization using active/decoy screening. Right: ROC AUC (x-axis) of receptor conformations plotted against enrichment factor (y-axis) and weighted enrichment factor (z-axis). Receptor conformations obtained from NMR models are shown with a circle marker, and clustered GaMD conformers are shown with a triangle marker. Left: ROC curve for the best performing conformer (1LXF NMR model 6) where the TPR is plotted against the FPR. The inset region shows the TPR and FPR of the top 40 docked compounds, the calculated EF, EF_weighted_, and highlights true positives.

### Exploration of Diphenyl Motifs Leads to Identification of Three cNTnC Calcium Sensitizers

Based on our previous screenings, we theorized lipophilic phenyl rings to be an important functional group for ligand binding into the cNTnC hydrophobic patch. Therefore, we focused our efforts on identification of small molecules containing diphenyl structures. We created three motifs of interest: diphenyl-imidazole (based on 2-[(4,5-diphenyl-1H-imidazol-2-yl)thio]-N-1,3-thiazol-2-ylacetamide), diphenyl-triazine, and diphenyl-methane (similar to 3-mDPA). The two-dimensional structures of these motifs are shown in **Figure S2**. We focused on two small molecule libraries from ChemBridge, the EXPRESS-Pick Library and Core Library, and filtered each library with a cheminformatics approach as described in the Methods section. For each compound of the filtered libraries (619,592 compounds) we calculated the Tanimoto coefficient to each of the three diphenyl motifs. Compounds found to have a high structural similarity (Tanimoto coefficient ≥ 0.6) were further filtered to remove redundancies from previous testing by our group and based on averaged Glide SP docking score across the three most predictive receptor conformations. This method led to the identification of 23 compounds for further experimental testing. Additionally, we performed a blind virtual screening of the filtered Core Library (412,866 compounds) and rank ordered compounds based on averaged Z-Score across the three most predictive receptor conformations. Upon visual inspection of the top 40 compounds, 17 were identified to have a diphenyl motif and were selected for further testing. In total, we ordered 40 compounds from ChemBridge for *in vitro* testing, the two-dimensional structures of all ordered compounds are available in **Figure S5**.

Initial stopped-flow kinetics revealed that 39 of the initial 40 compounds were soluble enough in aqueous buffer that experimental measurements could be obtained. Of the 39 compounds that were able to be experimentally tested, three showed at least a 15% decrease of the Ca^2+^ dissociation rate for the chimera. The average Ca^2+^ dissociation rate observed for the chimera in the absence of compounds was 68.6 ± 0.4 s^−1^. Compounds 2 (ChemBridge ID 7874460), 16 (ChemBridge ID 56598339), and 17 (ChemBridge ID 14233019) lead to a moderate slowing of the Ca^2+^ dissociation rate to 51.8 ± 0.4 *s*^−1^, 49.0 ± 0.3 *s*^−1^, and 57.6 ± 0.9 *s*^−1^ at 50 *μM*, respectively. To further characterize these hits, for each compound we performed stopped flow experiments using increasing concentrations of the respective small molecule in order to get a dose response. **Figure 4** illustrates the effects of each compound slowing the rate of Ca^2+^ disassociation from the chimera, where panel **4A** depicts the dose response and panel **4B** depicts a representative stopped-flow trace for each of the hit compounds at a 100 *μM* concentration of the compound. Of the three compounds, compound 17 was determined to have the lowest apparent experimental affinity of 482 ± 300 *μM*, a similar range to that of bepridil and NSC147866. Compound 2 was found to have an improved apparent experimental affinity (154 ± 23 *μM*) compared to compound 17. Compound 16 performed the best of the tested compounds with an apparent experimental affinity of 117 ± 17 *μM*. Compounds 2 and 16 had comparable affinities to known modulators TFP and W7, and showed significant improvement over levosimendan, bepridil, dfbp-o, and NSC147866. While we did not identify additional high affinity compounds, the three compounds that were identified meaningfully expand our knowledge of moderate affinity cNTnC binders.

**Figure 4.**
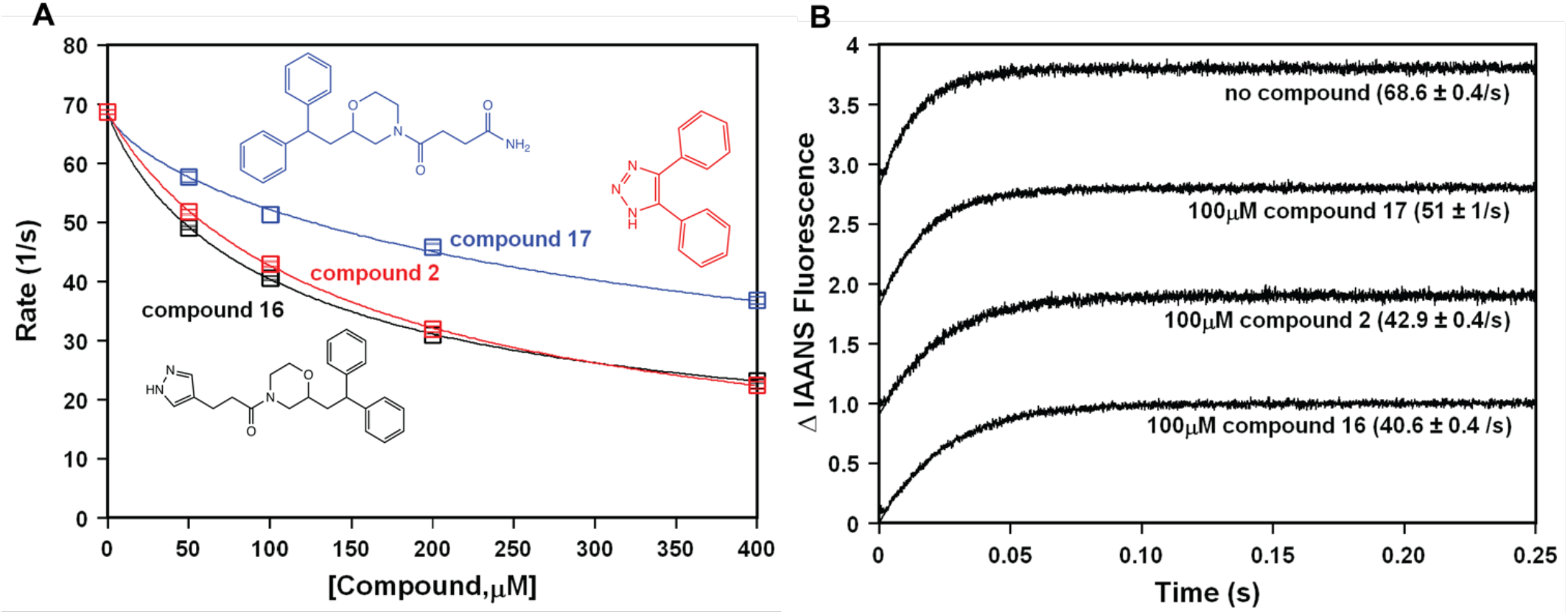
Effect of the hit compounds on the Ca^2+^ dissociation rate from the chimera. Panel (A) shows the plot of the apparent rates of Ca^2+^ dissociation from the chimera in the presence of increased concentration of compounds 2, 16 or 17. Each data point represents an average of at least three measurements ± standard error. Data were fit with an asymmetric sigmoid curve. Panel (B) shows representative stopped-flow traces as Ca^2+^ is removed from the chimera in the absence or presence of 100 μM of compounds 2, 16 or 17. The traces have been normalized and staggered for clarity.

Of the 40 tested compounds only five (12.5%) contained a diphenyl-imidazole structure and of these, one compound was identified to have moderate binding affinity: compound 2. Meanwhile 18 compounds (45%) contained diphenyl-triazine structures, which consistently failed to slow the rate of Ca^2+^ dissociation despite several compounds with a near-identical structure to previously identified modulators. We believe the substitution of a six-member ring disrupts the electrostatic and van der Waals protein-ligand interactions, thereby preventing the small molecule to bind deeply in the hydrophobic patch. Compounds containing diphenylmethane moieties accounted for 42.5% (17 compounds) of the screened compounds, and two of those small molecules were identified as medium to low affinity binders: compounds 16 and 17. Based on these results, we believe compounds containing a five-member ring (i.e., imidazole or triazole) or diphenylmethane are optimal for maximal lipophilic protein-ligand contact. On the other hand, our results seem to suggest that six-member rings (i.e., triazines) alter the sterics of the ligand and prevent deeper binding in the hydrophobic pocket. Representative docked poses of the two identified Ca^2+^ sensitizers with moderate affinity in this study are shown in **Figure 5**. In this figure, high affinity Ca^2+^ sensitizer 2-[(4,5-diphenyl-1H-imidazol-2-yl)thio]-N-1,3-thiazol-2-ylacetamide was docked into the cNTnC hydrophobic patch with the results of our virtual screenings overlayed, where panel **5A** shows compound 16 (ChemBridge ID 56598339) and panel **5B** shows compound 2 (ChemBridge ID 7874460). Compounds 2 and 16 were shown to have the prioritized diphenyl ring structure dock similarly to 2-[(4,5-diphenyl-1H-imidazol-2-yl)thio]-N-1,3-thiazol-2-ylacetamide. To confirm this, we determined the symmetry-corrected heavy atom RMSD of the diphenyl rings (utilizing spyrmsd ^88^) between 2-[(4,5-diphenyl-1H-imidazol-2-yl)thio]-N-1,3-thiazol-2-ylacetamide and compounds 2 and 16 to be 0.64 Å and 0.91 Å, respectively. While the diphenyl structural contacts of focus were maintained, it is likely that compounds 2 and 16 are missing other important protein-ligand interactions beyond the lipophilic structure. Therefore, future work should consider only five-member heterocycles or diphenylmethanes with a focus on optimizing ligand tail interactions with the cTnI_sp_.

**Figure 5.**
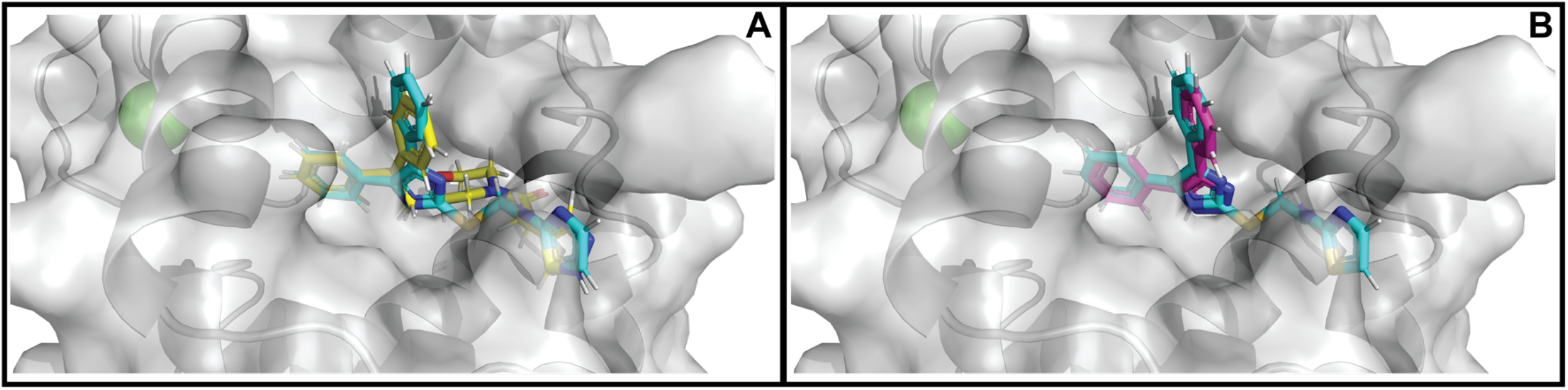
Docked poses of Ca^2+^ modulators in cNTnC hydrophobic pocket. PDB 1LXF model 6 is shown in gray surface representation with the protein backbone explicitly shown in cartoon representation, calcium sensitizer 2-[(4,5-diphenyl-1H-imidazol-2-yl)thio]-N-1,3-thiazol-2-ylacetamide shown in cyan overlayed with compound 16 shown in yellow (panel **A**) and compound 2 shown in magenta (panel **B**).

## Conclusion

In this study we tested one of our previously identified cTnC Ca^2+^ sensitizers (2-[(4,5-diphenyl-1H-imidazol-2-yl)thio]-N-1,3-thiazol-2-ylacetamide) for Ca^2+^ sensitization properties in the context of two different muscle types, cardiac trabeculae and slow skeletal muscle fibers which share the expression of cTnC and explored further virtual screening around a diphenyl motif. The results demonstrate that compound 7930079 is a potent sensitizer of force generation in cardiac trabeculae and in slow skeletal muscle fibers, with a much greater (four-fold) effect in the latter. This demonstrates the potential of this compound as a possible therapeutic for the treatment of heart failure and/or muscle weakness that is commonly associated with several myopathies, including cancer cachexia, AIDS and sarcopenia associated with aging.

We examined the predictive utility of GaMD clustered models based on NMR initial starting structures and, at least for cTnC, found no improvement compared to receptor conformers derived directly from NMR experiments. Future work should focus on whether this trend can be generalized across different receptor systems. Additionally, whether starting GaMD simulations from the most predictive models of 1LXF might increase the AUC and EF could be explored.

Through further drug discovery efforts centered around diphenyl motifs, we identified additional low affinity binders that increase Ca^2+^ sensitivity in cNTnC site II. While our approach did not lead to the desired success of identifying additional high affinity Ca^2+^ modulators, the observed binding affinities for two of the three identified compounds in this work were comparable to TFP, and better than levosimendan, W7, dfbp-o, bepridil, and NCI compound NSC147866. Therefore, we believe that the diphenyl-imidazole functional group is an important scaffold in cNTnC binding, but in itself is insufficient for high affinity binding. Future work to improve upon 2-[(4,5-diphenyl-1H-imidazol-2-yl)thio]-N-1,3-thiazol-2-ylacetamide will require collaboration with medicinal chemists. Lead optimization efforts will be focused on optimizing the protein-ligand interactions of the “tail” region, as this region was not sampled sufficiently in our present off the shelf catalog-based optimization. The identification of three more cNTnC binders increases our knowledge of the structure activity relationship in cTnC binding, which will be useful for follow up work.

## Acknowledgements

The authors would like to thank the members of the Lindert group for useful discussions. We would like to thank the Ohio Supercomputer Center for valuable computational resources ^89^. This work was supported by the NIH (R01 HL137015 to S.L.).

## Notes

### Competing Interest Statement

The authors have declared no competing interest.

