## Supplemental Tables and Figures for "Targeting Troponin C with Small Molecules Containing Diphenyl Moieties: Calcium Sensitivity Effects on Striated Muscle and Structure Activity Relationship"

Eric R. Hantz[1], Svetlana B. Tikunova[2], Natalya Belevych[3], Jonathan P. Davis[2], Peter J. Reiser[3, §], Steffen Lindert[1, §, *]
[1]Department of Chemistry and Biochemistry, The Ohio State University, Columbus, OH, 43210
[2]Davis Heart and Lung Research Institute and Department of Physiology and Cell Biology, The Ohio State University, Columbus, OH, 43210
[3]Division of Biosciences, College of Dentistry, The Ohio State University, Columbus, OH 43210

§ denotes co-senior authors

* Correspondence to:
Department of Chemistry and Biochemistry, Ohio State University
2114 Newman & Wolfrom Laboratory, 100 W. 18th Avenue, Columbus, OH 43210
614-292-8284 (office), 614-292-1685 (fax)


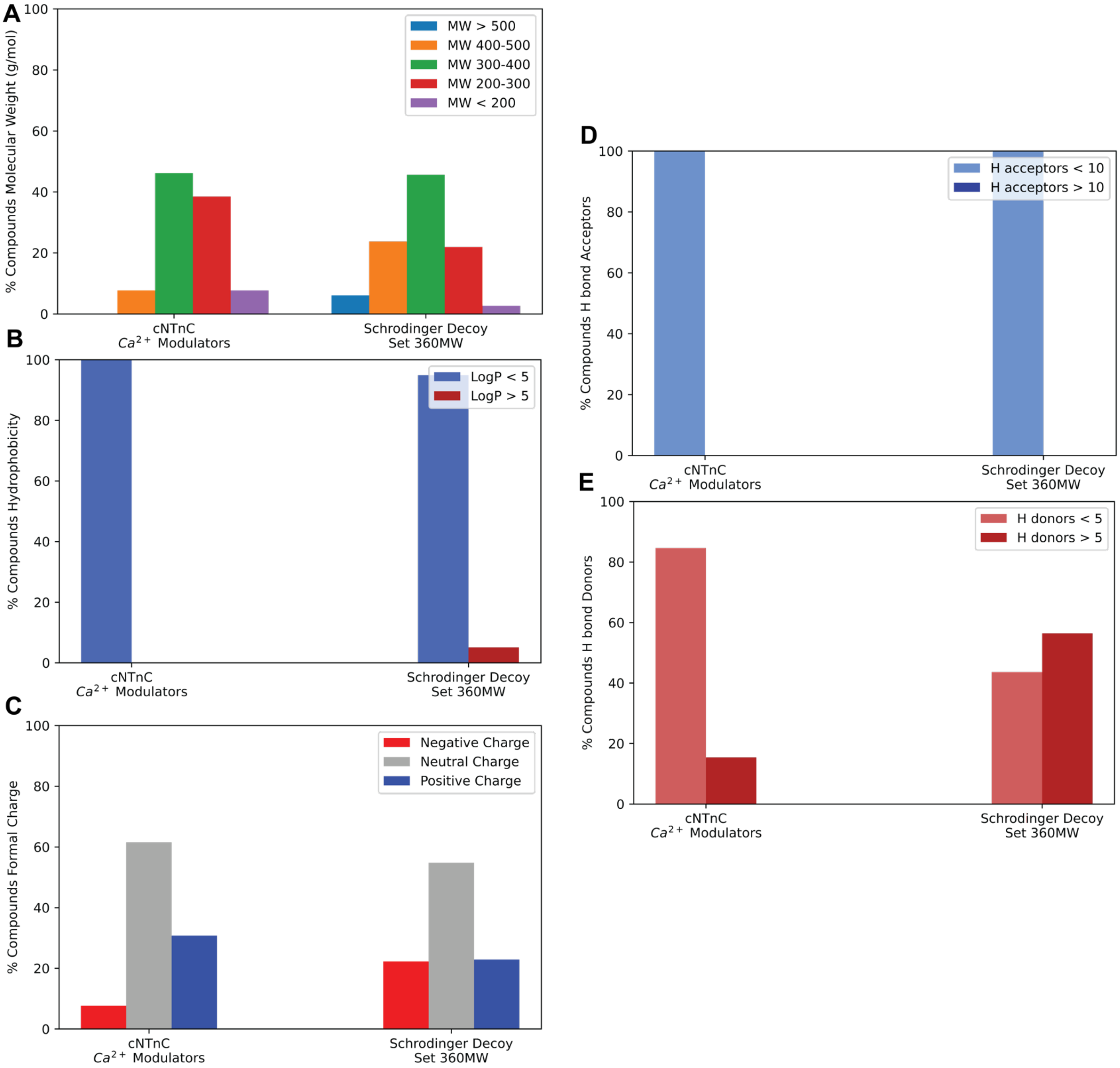


**Figure S1**. Comparison of various chemical properties between the 13 known cNTnC $Ca^{2+}$ modulators and Schrödinger decoy set with an average molecular weight of 360 g/mol. The properties shown are **(a)** Molecular weight (g/mol), **(b)** calculated LogP (hydrophobicity), **(c)** formal charge, **(d)** number of hydrogen bond acceptors, and **(e)** number of hydrogen bond donors.

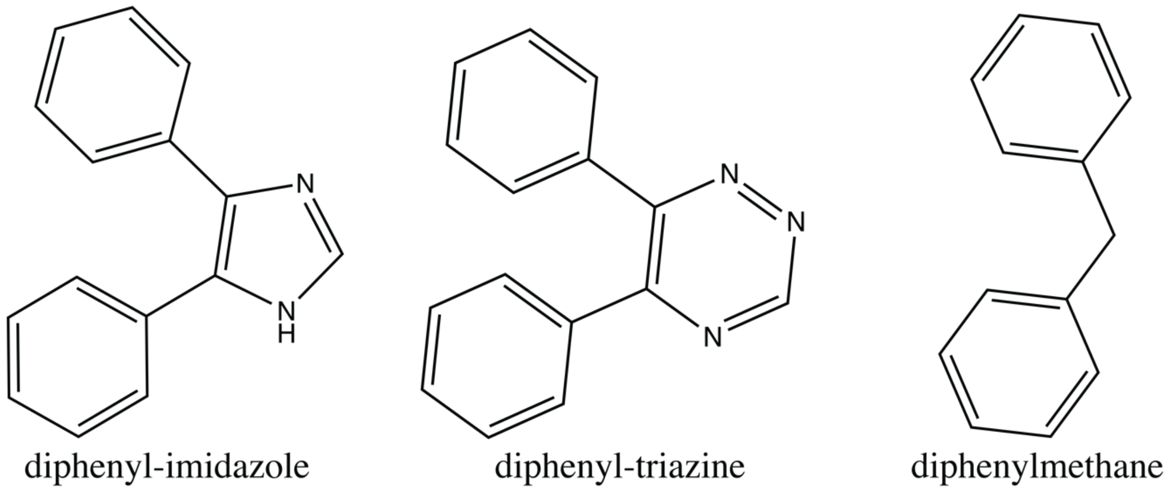


**Figure S2.** 2D structures of diphenyl motifs.

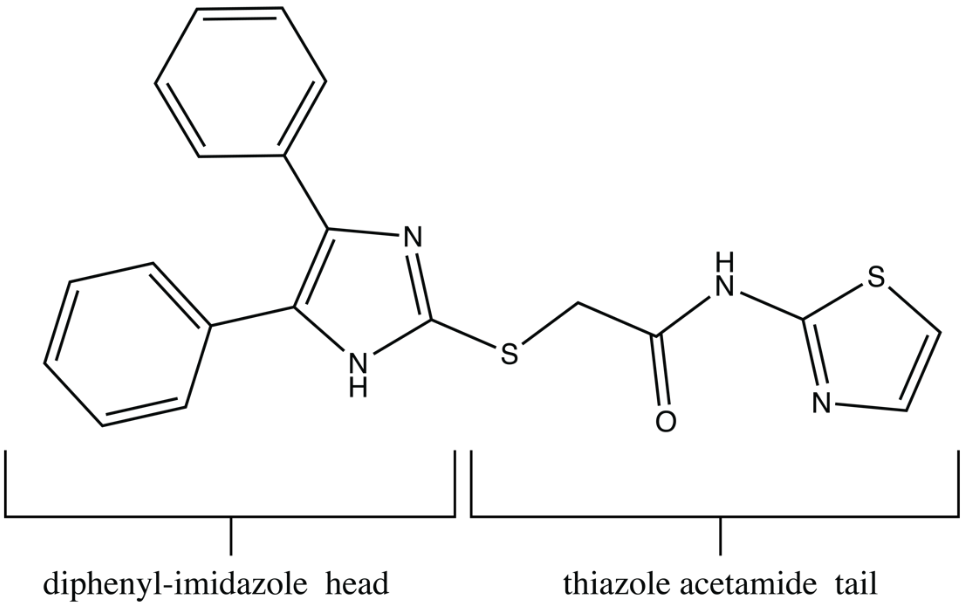


**Figure S3.** Naming convention for 2-[(4,5-diphenyl-1H-imidazol-2-yl)thio]-N-1,3-thiazol-2-ylacetamide (ChemBridge ID 7930079).

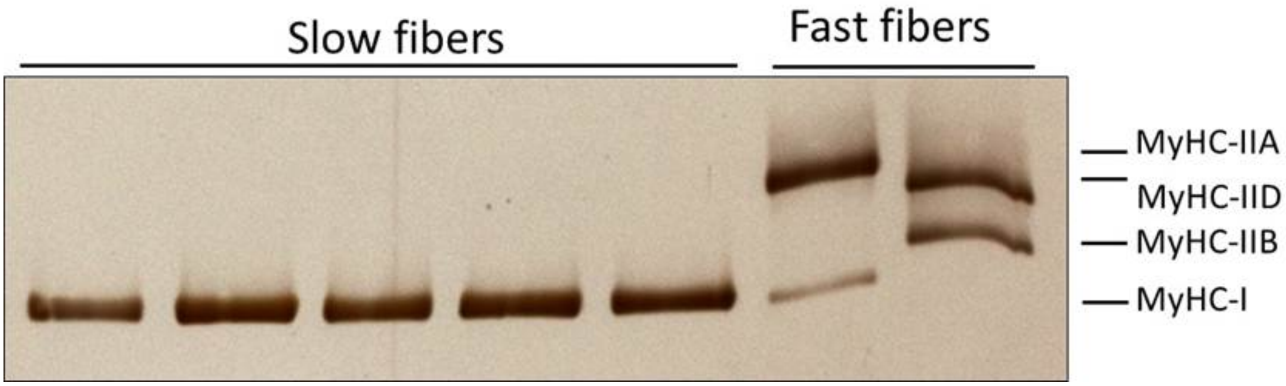


**Figure S4**. Myosin heavy chain (MyHC) isoforms in rat slow and fast skeletal muscle fibers. The image shows an example of the MyHC region of silver-stained gels that were loaded with single fibers from the slow soleus muscle and fast tibialis anterior muscle. The slow fibers were used for force/pCa measurements, and the fast fibers were used as sources of fast-type MyHC isoforms to distinguish the latter from MyHC-I. All ten of the studied soleus fibers expressed only slow-type MyHC-I. Fast-type MyHC-IIA, MyHC-IID and MyHC-IIB were expressed in the tibialis fibers on this gel. The first tibialis fiber (second last lane) was a fast/slow hybrid with MyHC-IIA and MyHC-I.

**Table S1**. Evaluation Metrics for all NMR Receptor Conformations

| PDB | Model Number | ROC AUC | EF | $EF_{weighted}$ |
|---|---|---|---|---|
| 1LXF | 1 | 0.676 | 7.79 | 5.98 |
| | 2 | 0.757 | 7.79 | 5.98 |
| | 3 | 0.730 | 3.90 | 4.49 |
| | 4 | 0.834 | 11.69 | 10.47 |
| | 5 | 0.812 | 7.79 | 8.97 |
| | 6 | 0.864 | 15.58 | 16.45 |
| | 7 | 0.782 | 7.79 | 10.47 |
| | 8 | 0.705 | 7.79 | 8.97 |
| | 9 | 0.517 | 1.95 | 1.50 |
| | 10 | 0.777 | 5.84 | 8.97 |
| | 11 | 0.780 | 7.79 | 7.48 |
| | 12 | 0.748 | 5.84 | 7.48 |
| | 13 | 0.647 | 7.79 | 7.48 |
| | 14 | 0.802 | 9.74 | 11.96 |
| | 15 | 0.786 | 7.79 | 5.98 |
| | 16 | 0.767 | 11.69 | 11.96 |
| | 17 | 0.754 | 9.74 | 10.47 |
| | 18 | 0.682 | 7.79 | 10.47 |
| | 19 | 0.691 | 7.79 | 8.97 |
| | 20 | 0.748 | 1.95 | 1.50 |
| | 21 | 0.756 | 11.69 | 11.96 |
| | 22 | 0.788 | 7.79 | 7.48 |
| | 23 | 0.828 | 13.64 | 14.96 |
| | 24 | 0.631 | 1.95 | 2.99 |
| | 25 | 0.791 | 7.79 | 7.48 |
| | 26 | 0.792 | 5.84 | 5.98 |
| | 27 | 0.797 | 7.79 | 5.98 |
| | 28 | 0.858 | 13.64 | 14.96 |
| | 29 | 0.857 | 11.69 | 11.96 |
| | 30 | 0.716 | 1.95 | 2.99 |
| 1MXL | 1 | 0.583 | 1.95 | 2.99 |
| | 2 | 0.577 | 0.00 | 0.00 |
| | 3 | 0.523 | 1.95 | 1.50 |
| | 4 | 0.383 | 0.00 | 0.00 |
| | 5 | 0.429 | 0.00 | 0.00 |
| | 6 | 0.531 | 0.00 | 0.00 |
| | 7 | 0.518 | 1.95 | 1.50 |
| | 8 | 0.421 | 1.95 | 1.50 |
| | 9 | 0.529 | 0.00 | 0.00 |
| | 10 | 0.496 | 1.95 | 1.50 |
| | 11 | 0.532 | 1.95 | 1.50 |
| | 12 | 0.210 | 0.00 | 0.00 |
| | 13 | 0.697 | 1.95 | 2.99 |

| PDB | Model Number | ROC AUC | EF | $EF_{weighted}$ |
|---|---|---|---|---|
| 1MXL | 14 | 0.499 | 0.00 | 0.00 |
| | 15 | 0.402 | 3.90 | 2.99 |
| | 16 | 0.554 | 0.00 | 0.00 |
| | 17 | 0.213 | 1.95 | 1.50 |
| | 18 | 0.470 | 1.95 | 1.50 |
| | 19 | 0.323 | 1.95 | 1.50 |
| | 20 | 0.177 | 0.00 | 0.00 |
| | 21 | 0.556 | 0.00 | 0.00 |
| | 22 | 0.704 | 0.00 | 0.00 |
| | 23 | 0.701 | 1.95 | 2.99 |
| | 24 | 0.562 | 0.00 | 0.00 |
| | 25 | 0.538 | 0.00 | 0.00 |
| | 26 | 0.162 | 1.95 | 1.50 |
| | 27 | 0.745 | 5.84 | 4.49 |
| | 28 | 0.139 | 0.00 | 0.00 |
| | 29 | 0.654 | 0.00 | 0.00 |
| | 30 | 0.500 | 3.90 | 2.99 |
| | 31 | 0.499 | 0.00 | 0.00 |
| | 32 | 0.710 | 1.95 | 2.99 |
| | 33 | 0.485 | 3.90 | 2.99 |
| | 34 | 0.687 | 1.95 | 2.99 |
| | 35 | 0.288 | 0.00 | 0.00 |
| | 36 | 0.182 | 0.00 | 0.00 |
| | 37 | 0.560 | 1.95 | 2.99 |
| | 38 | 0.540 | 0.00 | 0.00 |
| | 39 | 0.472 | 3.90 | 2.99 |
| | 40 | 0.529 | 1.95 | 2.99 |
| 2KFX | 1 | 0.699 | 0.00 | 0.00 |
| | 2 | 0.631 | 3.90 | 2.99 |
| | 3 | 0.695 | 1.95 | 1.50 |
| | 4 | 0.719 | 3.90 | 2.99 |
| | 5 | 0.636 | 0.00 | 0.00 |
| | 6 | 0.683 | 3.90 | 2.99 |
| | 7 | 0.715 | 0.00 | 0.00 |
| | 8 | 0.693 | 1.95 | 1.50 |
| | 9 | 0.706 | 0.00 | 0.00 |
| | 10 | 0.761 | 3.90 | 4.49 |
| 2KRD | 1 | 0.702 | 1.93 | 2.99 |
| | 2 | 0.807 | 5.84 | 4.49 |
| | 3 | 0.736 | 3.90 | 2.99 |
| | 4 | 0.738 | 1.95 | 1.50 |
| | 5 | 0.683 | 1.95 | 1.50 |
| | 6 | 0.646 | 3.90 | 2.99 |
| | 7 | 0.727 | 7.79 | 8.97 |

| PDB | Model Number | ROC AUC | EF | $EF_{weighted}$ |
|---|---|---|---|---|
| 2KRD | 8 | 0.670 | 3.90 | 2.99 |
| | 9 | 0.714 | 3.90 | 4.49 |
| | 10 | 0.737 | 0.00 | 0.00 |
| | 11 | 0.744 | 3.90 | 2.99 |
| | 12 | 0.821 | 5.84 | 4.49 |
| | 13 | 0.720 | 0.00 | 0.00 |
| | 14 | 0.698 | 0.00 | 0.00 |
| | 15 | 0.716 | 3.90 | 2.99 |
| | 16 | 0.632 | 3.90 | 2.99 |
| | 17 | 0.807 | 7.79 | 7.48 |
| | 18 | 0.713 | 5.84 | 4.49 |
| | 19 | 0.649 | 0.00 | 0.00 |
| | 20 | 0.627 | 0.00 | 0.00 |
| 2L1R | 1 | 0.717 | 5.84 | 5.98 |
| | 2 | 0.565 | 1.95 | 1.50 |
| | 3 | 0.623 | 5.84 | 4.49 |
| | 4 | 0.484 | 1.95 | 1.50 |
| | 5 | 0.565 | 3.90 | 2.99 |
| | 6 | 0.674 | 3.90 | 2.99 |
| | 7 | 0.585 | 0.00 | 0.00 |
| | 8 | 0.794 | 5.84 | 5.98 |
| | 9 | 0.736 | 3.90 | 2.99 |
| | 10 | 0.554 | 3.90 | 2.99 |
| | 11 | 0.508 | 3.90 | 4.49 |
| | 12 | 0.742 | 7.79 | 10.47 |
| | 13 | 0.653 | 3.90 | 2.99 |
| | 14 | 0.577 | 0.00 | 0.00 |
| | 15 | 0.734 | 5.84 | 4.49 |
| | 16 | 0.602 | 3.90 | 2.99 |
| | 17 | 0.580 | 0.00 | 0.00 |
| | 18 | 0.657 | 3.90 | 2.99 |
| | 19 | 0.662 | 0.00 | 0.00 |
| | 20 | 0.559 | 3.90 | 2.99 |
| 2MKP | 1 | 0.110 | 0.00 | 0.00 |
| | 2 | 0.517 | 0.00 | 0.00 |
| | 3 | 0.437 | 1.95 | 1.50 |
| | 4 | 0.365 | 0.00 | 0.00 |
| | 5 | 0.152 | 3.90 | 2.99 |
| | 6 | 0.188 | 0.00 | 0.00 |
| | 7 | 0.215 | 1.95 | 1.50 |
| | 8 | 0.406 | 0.00 | 0.00 |
| | 9 | 0.555 | 0.00 | 0.00 |
| | 10 | 0.654 | 0.00 | 0.00 |
| 5W88 | 1 | 0.205 | 0.00 | 0.00 |

| PDB | Model Number | ROC AUC | EF | $EF_{weighted}$ |
|---|---|---|---|---|
| 5W88 | 2 | 0.532 | 3.90 | 2.99 |
| | 3 | 0.212 | 0.00 | 0.00 |
| | 4 | 0.635 | 0.00 | 0.00 |
| | 5 | 0.145 | 1.95 | 1.50 |
| | 6 | 0.071 | 0.00 | 0.00 |
| | 7 | 0.211 | 0.00 | 0.00 |
| | 8 | 0.218 | 3.90 | 4.49 |
| 5WCL | 1 | 0.483 | 3.90 | 2.99 |
| | 2 | 0.484 | 3.90 | 2.99 |
| | 3 | 0.591 | 1.95 | 1.50 |
| | 4 | 0.533 | 3.90 | 2.99 |
| | 5 | 0.441 | 3.90 | 2.99 |
| | 6 | 0.653 | 5.84 | 5.98 |
| | 7 | 0.599 | 5.84 | 4.49 |
| | 8 | 0.518 | 0.00 | 0.00 |
| 6MV3 | 1 | 0.466 | 1.95 | 1.50 |
| | 2 | 0.700 | 3.90 | 4.49 |
| | 3 | 0.520 | 1.95 | 1.50 |
| | 4 | 0.632 | 1.95 | 1.50 |
| | 5 | 0.520 | 0.00 | 0.00 |
| | 6 | 0.480 | 1.95 | 2.99 |
| | 7 | 0.657 | 3.90 | 2.99 |
| | 8 | 0.499 | 0.00 | 0.00 |
| | 9 | 0.636 | 0.00 | 0.00 |
| | 10 | 0.618 | 1.95 | 1.50 |
| | 11 | 0.532 | 1.95 | 1.50 |
| | 12 | 0.610 | 1.95 | 1.50 |
| | 13 | 0.466 | 0.00 | 0.00 |
| | 14 | 0.617 | 1.95 | 1.50 |
| | 15 | 0.553 | 1.95 | 1.50 |
| | 16 | 0.571 | 3.90 | 2.99 |
| | 17 | 0.655 | 1.95 | 1.50 |
| | 18 | 0.658 | 1.95 | 1.50 |
| | 19 | 0.658 | 1.95 | 1.50 |
| | 20 | 0.629 | 0.00 | 0.00 |

**Table S2.** Evaluation Metrics for all GaMD Generated Receptor Conformers

| PDB | Clustered Model Number | ROC AUC | EF | $EF_{weighted}$ |
|---|---|---|---|---|
| 1LXF | 0 | 0.649 | 5.84 | 5.98 |
| | 1 | 0.739 | 7.79 | 8.97 |
| | 2 | 0.748 | 5.84 | 8.97 |
| | 3 | 0.603 | 3.90 | 5.98 |
| | 4 | 0.626 | 1.95 | 1.50 |
| | 5 | 0.695 | 3.90 | 5.98 |
| | 6 | 0.678 | 1.95 | 2.99 |
| 1MXL | 0 | 0.292 | 0.00 | 0.00 |
| | 1 | 0.339 | 1.95 | 2.99 |
| | 2 | 0.607 | 1.95 | 1.50 |
| | 3 | 0.495 | 0.00 | 0.00 |
| | 4 | 0.472 | 0.00 | 0.00 |
| | 5 | 0.000 | 0.00 | 0.00 |
| | 6 | 0.384 | 0.00 | 0.00 |
| | 7 | 0.310 | 0.00 | 0.00 |
| | 8 | 0.539 | 1.95 | 1.50 |
| | 9 | 0.000 | 0.00 | 0.00 |
| | 10 | 0.533 | 0.00 | 0.00 |
| | 11 | 0.212 | 1.95 | 2.99 |
| | 12 | 0.585 | 1.95 | 2.99 |
| | 13 | 0.390 | 0.00 | 0.00 |
| 2KFX | 0 | 0.225 | 0.00 | 0.00 |
| | 1 | 0.657 | 3.90 | 2.99 |
| | 2 | 0.539 | 1.95 | 1.50 |
| | 3 | 0.657 | 5.84 | 4.49 |
| | 4 | 0.739 | 1.95 | 1.50 |
| | 5 | 0.686 | 3.90 | 4.49 |
| | 6 | 0.698 | 0.00 | 0.00 |
| 2KRD | 0 | 0.682 | 1.95 | 1.50 |
| | 1 | 0.621 | 0.00 | 0.00 |
| | 2 | 0.466 | 5.84 | 4.49 |
| | 3 | 0.378 | 0.00 | 0.00 |
| | 4 | 0.515 | 0.00 | 0.00 |
| | 5 | 0.643 | 3.90 | 4.49 |
| | 6 | 0.504 | 1.95 | 1.50 |
| | 7 | 0.730 | 1.95 | 1.50 |
| | 8 | 0.569 | 0.00 | 0.00 |
| | 9 | 0.497 | 0.00 | 0.00 |
| | 10 | 0.546 | 0.00 | 0.00 |
| | 11 | 0.437 | 3.90 | 2.99 |
| 2L1R | 0 | 0.320 | 0.00 | 0.00 |
| | 1 | 0.276 | 0.00 | 0.00 |
| | 2 | 0.137 | 0.00 | 0.00 |

| PDB | Clustered Model Number | ROC AUC | EF | $EF_{weighted}$ |
|---|---|---|---|---|
| 2L1R | 3 | 0.367 | 0.00 | 0.00 |
| | 4 | 0.075 | 1.95 | 1.50 |
| | 5 | 0.381 | 1.95 | 1.50 |
| | 6 | 0.000 | 0.00 | 0.00 |
| | 7 | 0.078 | 1.95 | 1.50 |
| | 8 | 0.249 | 0.00 | 0.00 |
| | 9 | 0.131 | 0.00 | 0.00 |
| 2MKP | 0 | 0.072 | 0.00 | 0.00 |
| | 1 | 0.208 | 0.00 | 0.00 |
| | 2 | 0.074 | 1.95 | 1.50 |
| | 3 | 0.412 | 0.00 | 0.00 |
| | 4 | 0.597 | 3.90 | 2.99 |
| | 5 | 0.519 | 1.95 | 1.50 |
| | 6 | 0.000 | 0.00 | 0.00 |
| | 7 | 0.399 | 1.95 | 1.50 |
| | 8 | 0.142 | 1.95 | 1.50 |
| | 9 | 0.689 | 0.00 | 0.00 |
| | 10 | 0.270 | 1.95 | 1.50 |
| | 11 | 0.000 | 0.00 | 0.00 |
| | 12 | 0.535 | 0.00 | 0.00 |
| | 13 | 0.307 | 0.00 | 0.00 |
| 5W88 | 0 | 0.279 | 5.84 | 4.49 |
| | 1 | 0.795 | 5.84 | 8.97 |
| | 2 | 0.449 | 0.00 | 0.00 |
| | 3 | 0.423 | 1.95 | 2.99 |
| | 4 | 0.750 | 3.90 | 5.98 |
| | 5 | 0.576 | 3.90 | 2.99 |
| | 6 | 0.654 | 5.84 | 4.49 |
| | 7 | 0.673 | 3.90 | 4.49 |
| | 8 | 0.584 | 7.79 | 5.98 |
| | 9 | 0.720 | 1.95 | 1.50 |
| | 10 | 0.661 | 1.95 | 2.99 |
| | 11 | 0.668 | 3.90 | 4.49 |
| | 12 | 0.688 | 1.95 | 1.50 |
| | 13 | 0.661 | 1.95 | 2.99 |
| 5WCL | 0 | 0.530 | 1.95 | 1.50 |
| | 1 | 0.420 | 3.90 | 4.49 |
| | 2 | 0.523 | 5.84 | 5.98 |
| | 3 | 0.544 | 7.79 | 5.98 |
| | 4 | 0.590 | 13.64 | 11.96 |
| | 5 | 0.542 | 5.84 | 4.49 |
| | 6 | 0.750 | 5.84 | 5.98 |
| | 7 | 0.539 | 3.90 | 2.99 |
| | 8 | 0.418 | 3.90 | 4.49 |

| PDB | Clustered Model Number | ROC AUC | EF | $EF_{weighted}$ |
|---|---|---|---|---|
| 5WCL | 9 | 0.504 | 7.79 | 7.48 |
| | 10 | 0.467 | 1.95 | 1.50 |
| 6MV3 | 0 | 0.484 | 0.00 | 0.00 |
| | 1 | 0.613 | 1.95 | 2.99 |
| | 2 | 0.513 | 0.00 | 0.00 |
| | 3 | 0.600 | 0.00 | 0.00 |
| | 4 | 0.483 | 0.00 | 0.00 |
| | 5 | 0.602 | 3.90 | 4.49 |
| | 6 | 0.471 | 0.00 | 0.00 |
| | 7 | 0.559 | 0.00 | 0.00 |
| | 8 | 0.514 | 0.00 | 0.00 |
| | 9 | 0.397 | 0.00 | 0.00 |
| | 10 | 0.456 | 1.95 | 2.99 |
| | 11 | 0.600 | 0.00 | 0.00 |

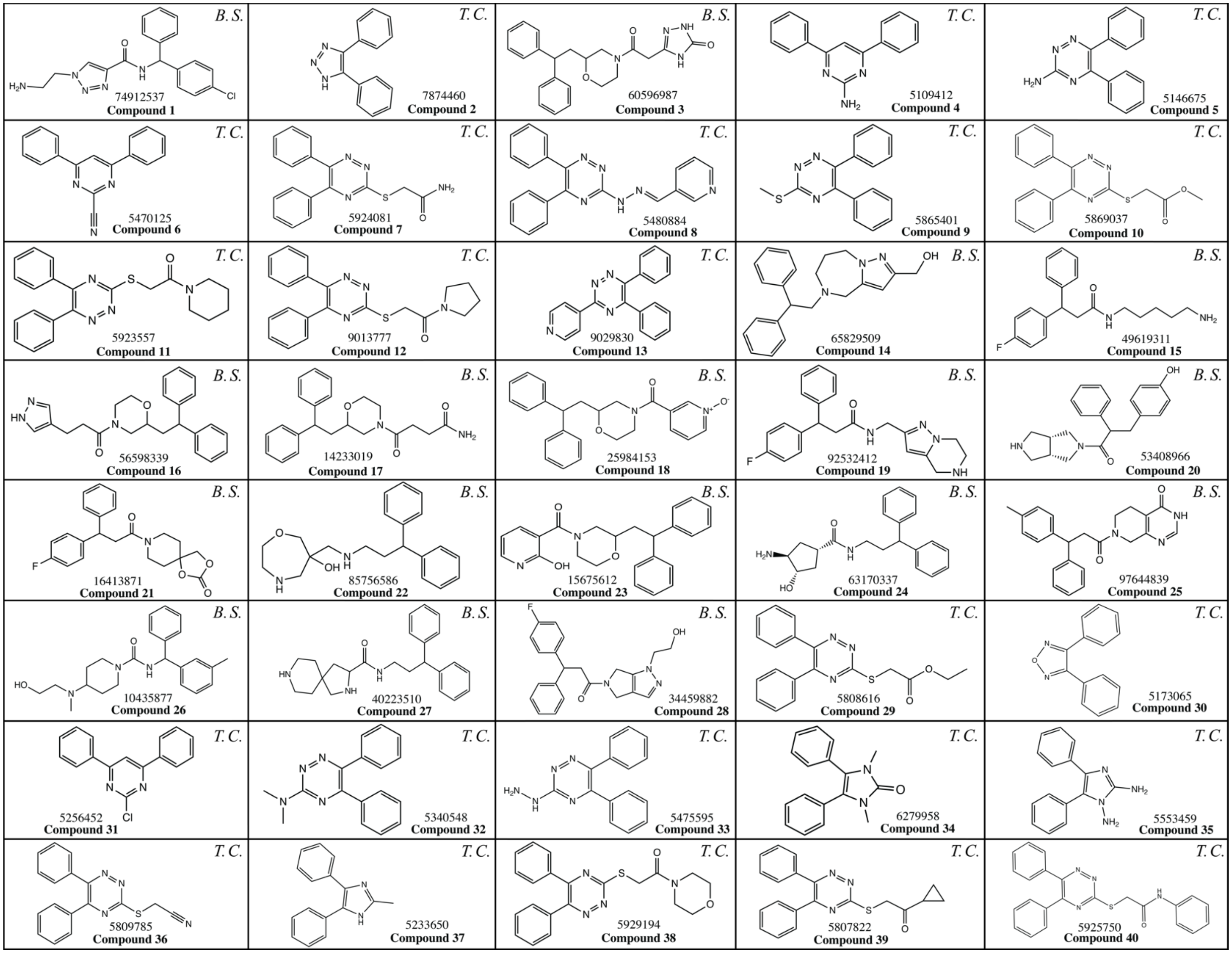


**Figure S5.** 2D structures of all ordered ChemBridge compounds. ChemBridge compound IDs and compounds numbered in order of *in vitro* testing (bolded). Compounds ordered based on Tanimoto coefficient (T.C.) or blind screening of ChemBridge Core library (B.S.) are designated in the top right corner of the compound's respective box.
